# The allosteric mechanism of G-protein-coupled receptors is induced fit, not conformational selection

**DOI:** 10.1101/2025.01.28.635241

**Authors:** Kazem Asadollahi, Paul R. Gooley, Thomas R. Weikl

**Affiliations:** Department of Biochemistry and Pharmacology, Bio21 Molecular Science and Biotechnology Institute, University of Melbourne, Parkville, VIC 3052, Australia; Department of Biomolecular Systems, Max Planck Institute of Colloids and Interfaces, 14476 Potsdam, Germany

## Abstract

The allosteric mechanism of G-protein-coupled receptors (GPCRs) involves a population shift from inactive to active receptor conformations in response to the binding of ligand agonists. Two possible kinetic mechanisms for this population shift are induced fit and conformational selection. The two mechanisms differ in the temporal sequence of binding events and conformational changes: Ligand bindings occurs prior to the change from the inactive to the active receptor conformation in the induced-fit mechanism, and after the conformational change in the conformational-selection mechanism. In this article, we discuss the current evidence from experiments that probe the binding kinetics of GPCRs to identify the allosteric mechanism. For the peptide-activated neurotensin receptor 1, the modeling of kinetic data from stopped-flow mixing experiments indicates an induced-fit mechanism ^1^. The conformational exchange rates of the induced-fit mechanism obtained from this modeling agree with rates measured by saturation transfer difference NMR experiments of the peptide-receptor complex, which corroborates the mechanism. For the small-molecule-activated β_2_-andrenergic receptor, an induced-fit mechanism has been inferred from a comparison of ligand-association rates for the inactive and the active receptor conformation ^2^. A stabilization of the active receptor conformation by G proteins or nanobodies leads to a decrease of ligand association rates, which indicates that ligand binding occurs in the inactive conformation and, thus, prior to the change from the inactive to the active conformation as in the induced-fit mechanism. A structural explanation for the induced-fit mechanism of the β_2_-andrenergic receptor is a closed lid over the binding site that blocks ligand entry in the active conformation. Since constriction and closing of the ligand-binding site in the active conformation is rather common for small-molecule-activated and peptide-activated GPCRs, induced fit likely is shared as allosteric mechanism by these GPCRs.

## Population shift as basis for GPCR allostery

G-protein-coupled receptors (GPCRs) constitute the largest super-family of cell membrane proteins and mediate the majority of cellular responses to extracellular stimuli ^3^. Binding of agonist ligands to the extracellular side of GPCRs induces global conformational changes in the seven transmembrane helix domain that lead to the activation of G proteins bound to the intracellular side of GPCRs.

The allosteric coupling of extracellular and intracellular binding sites of GPCRs can be understood from a chemical equilibrium of inactive and active (i.e. G-protein-activating) conformations that is shifted by ligand binding ^4,5^ (see Fig.1). In the unbound state, the inactive conformational ensemble R_1_ is predominantly populated, with a minor population of the active conformational ensemble R_2_ leading to a basal, ligand-independent level of activity. In the ligand-bound state, the chemical equilibrium is shifted towards the active conformational ensemble R_2_L.

## Different notions of conformational selection

The population shift in the equilibrium model of Fig. 1 is the basis for understanding GPCR allostery, and protein allostery in general ^7,8^. However, this equilibrium model does not state a kinetic sequence of events in GPCR activation. In general, two possible temporal sequences in the coupling of binding events and conformational changes of proteins are: The binding event occurs either prior to or after the conformational change ^9–12^. Such a clear temporal sequence of events implies that the dwell (or residence) times in the different conformational and binding states of a protein are much larger than the transition times between the states, so that a temporal overlap of conformational transitions and binding events is unlikely ^13^.

**Fig. 1.**
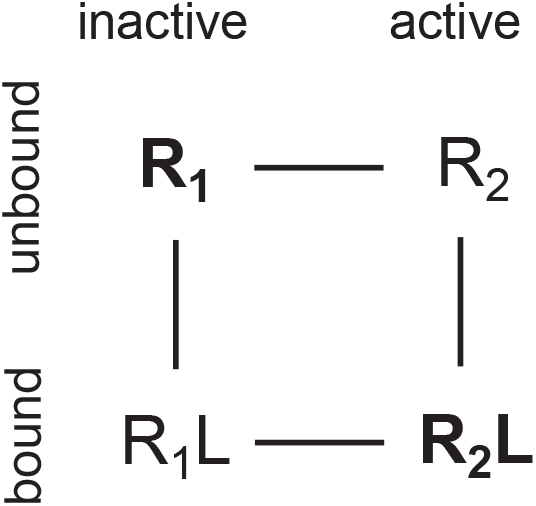
Population-shift equilibrium model of protein allostery. In the unbound state, the inactive conformational ensemble R_1_ of the protein is predominantly populated (indicated in bold), and the population of the active conformational ensemble R_2_ is minor. In the ligand-bound state, the populations are shifted, with a predominant population of R_2_L (indicated in bold) and a minor population of R_1_L.

A sequence of events in which ligand binding occurs prior to the conformational change of a protein has been termed induced fit ^14^. The opposing sequence of events in which ligand binding occurs after the conformational change of the protein is generally denoted as conformational selection ^15^. However, the term conformational selection has also been used in different or broader meanings ^16^. In the GPCR field, the term conformational selection has been used rather synonymously to the equilibrium concept of population shift as general basis for allostery ^4,17–20^.

## Induced fit or conformational selection?

The question of this article is: Does ligand-activation of GPCRs occur via induced fit or conformational selection? In this question, the term conformational selection is used for a kinetic mechanism. The population shift in the equilibrium model of Fig.1 is the basis for allostery, but we now ask: What is the kinetic mechanism, or the kinetic sequence of events that leads to this population shift?

Induced fit and conformational selection differ in the conformational ensemble in which ligand binding occurs (see Fig. 2). In the induced-fit mechanism, the ligand L binds to the inactive conformational ensemble R_1_. Along the induced-fit activation pathway from R_1_ to R_2_L, ligand binding therefore occurs prior to the conformational change. A conformational change from R_1_ to R_2_ can also occur in the unbound state, but this conformational change is “ off-pathway” to activation in the induced-fit mechanism. In the conformational-selection mechanism, in contrast, the ligand binds to the active conformational ensemble R_2_. In this mechanism, ligand binding therefore occurs prior to the conformational change along the activation pathway from R_1_ to R_2_L. In principle, the conformational-selection and induced-fit pathways could also be parallel, alternative pathways. In practice, however, one of these pathways is likely the dominant pathway at the relevant ligand concentrations ^21^, or the only possible pathway if there are structural reasons that prevent ligand binding to one of the conformational ensembles R_1_ or R_2_ ^22^.

**Fig. 2.**
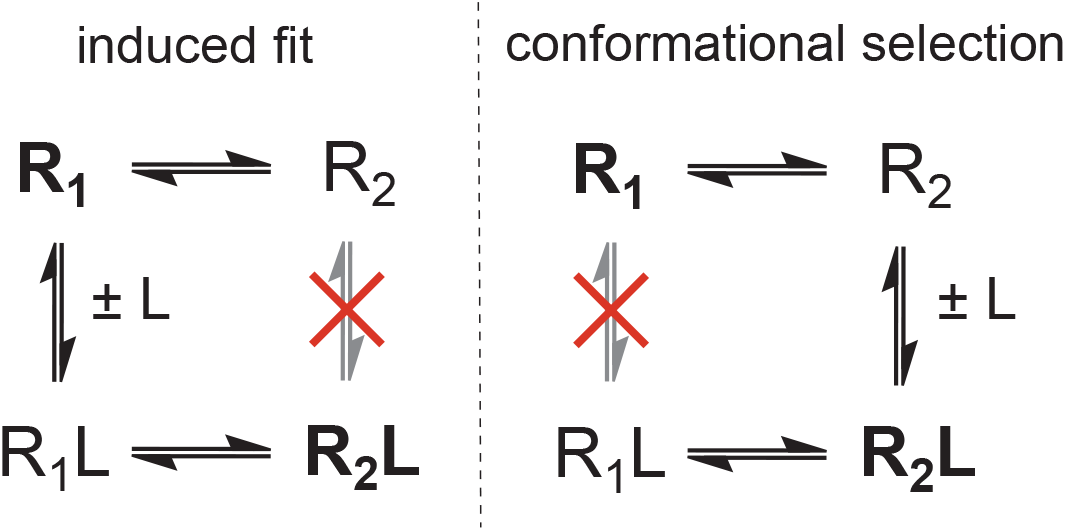
Kinetic models of protein allostery. In the induced-fit model, only the inactive conformational ensemble R_1_ of the protein is binding-competent, and ligand binding occurs prior to the conformational change of the protein on the activation pathway from R_1_ to R_2_L. In the conformational-selection model, only the active conformational ensemble R_2_ is binding-competent, and ligand binding occurs after the conformational change of the protein on the activation pathway from R_1_ to R_2_L.

Identifying the dominant binding mechanism requires to probe the binding kinetics. Thermodynamic principles dictate that equilibrium properties such as the equilibrium population of states are independent of the pathways along which the states can be reached. For example, the basal activity of a GPCR indicates that the active conformational ensemble R_2_ is populated to some extent in the unbound state of the GPCR. However, this does not allow to infer whether R_2_ is on-pathway (conformational selection) or off-pathway (induced fit) regarding the activation path from R_1_ to R_2_L. In other words, a minor population of R_2_ is a necessary condition for the conformational-selection binding pathway on which R_2_ occurs as intermediate, but is not a sufficient condition to infer conformational selection.

## Distinguishing induced fit and conformational selection with stopped-flow mixing experiments

Stopped-flow mixing experiments probe the chemical relaxation into binding equilibrium. For both induced-fit and conformational-selection binding processes, the final relaxation into equilibrium can be described as a double-exponential relaxation with two relaxation rates *k*_1_ and *k*_2_ ^6^. But the two binding mechanisms differ in how *k*_1_ and *k*_2_ depend on the ligand concentration [L]_0_ (see Fig. 3), which can be used to distinguish induced fit and conformational selection. For an induced-fit binding process, the functions *k*_1_([L]_0_) and *k*_2_([L]_0_) are symmetric around a minimum. For a conformational-selection process, in contrast, *k*_1_([L]_0_) and *k*_2_([L]_0_) do not exhibit this symmetry: the smaller rate *k*_2_ either decreases monotonously with [L]_0_ if the conformational excitation rate *k*_*e*_ of the process is smaller than the dissociation rate *k*_−_, or exhibits a minimum between asymmetric arms for *k*_*e*_ *> k*_−_ (see Fig. 3).

**Fig. 3.**
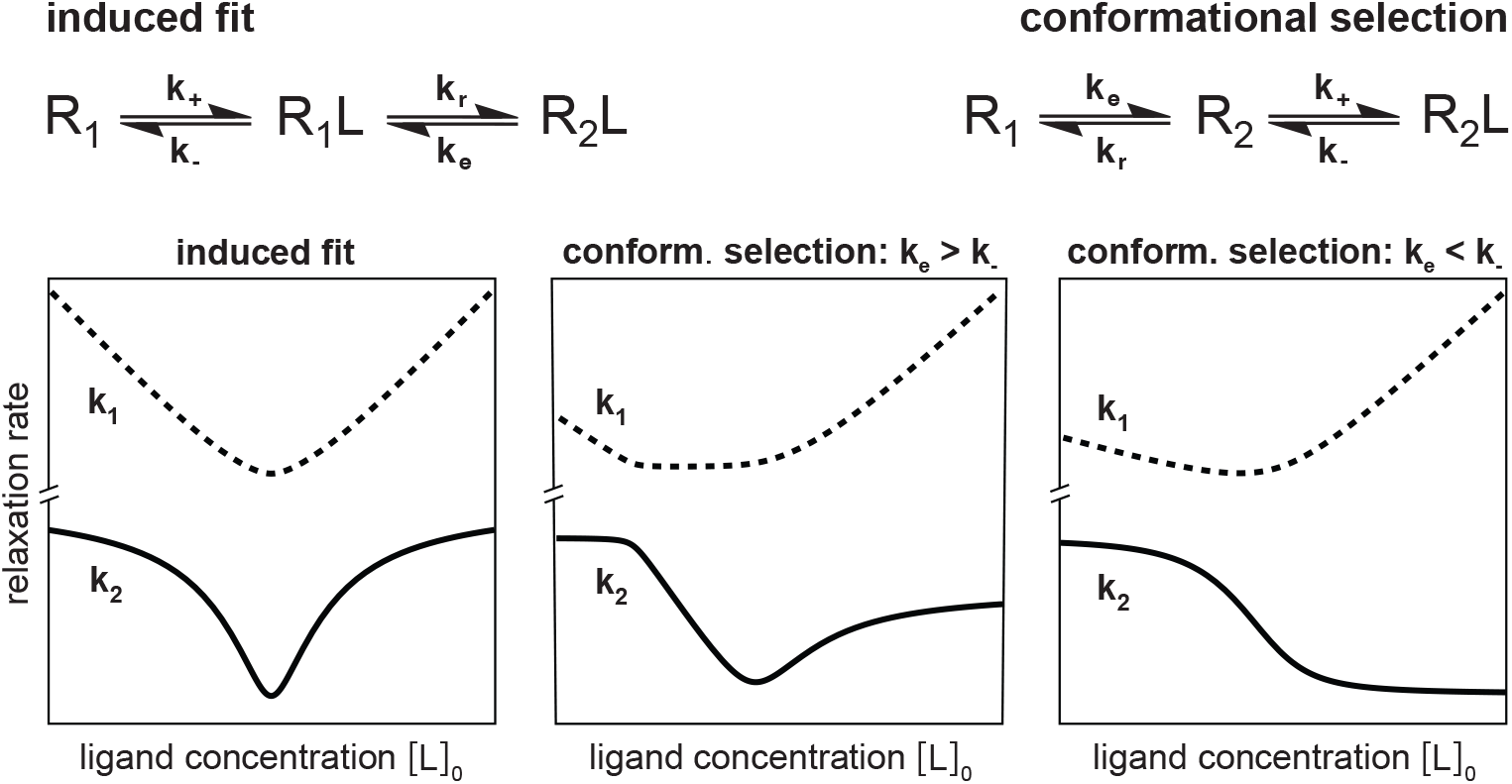
Characteristic relaxation rates *k*_1_ and *k*_2_ of the induced-fit and conformational-selection model as functions of the ligand concentration [L]_0_ in mixing experiments 6. In the induced-fit model, *k*_1_ and *k*_2_ are symmetric functions around a minimum located at 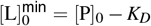 where [P]_0_ is the protein concentration in the mixture and *K*_*D*_ = *k*_−_*k*_*e*_*/*(*k*_+_(*k*_*e*_ + *k*_*r*_)) is the dissociation constant in the induced-fit model. In the conformational-selection model, the smaller relaxation rate *k*_2_, which dominates the final relaxation into equilibrium, monotonously decreases with [L]_0_ if the conformational excitation rate *k*_*e*_ for the transition from R_1_ to R_2_ is smaller than the dissociation rate *k*_−_ of the complex R_2_L. If *k*_*e*_ is larger than *k*_−_, the relaxation rate *k*_2_ exhibits a minimum at 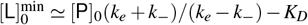 with dissociation constant *K*_*D*_ = *k*_−_(*k*_*e*_ + *k*_*r*_)*/*(*k*_+_*k*_*e*_) in the conformational-selection model, without the symmetry of the induced-fit model. The relaxation rates *k*_1_ and *k*_2_ here have been calculated for the 3-state pathways from R_1_ to R_2_L in induced fit and conformational selection, because the rates are not affected by the additional “ off-pathway” conformational change in the 4-state models of Fig. 2 if the sum of the rates for this exchange is larger than *k*_1_ and *k*_2_, which is plausible for a typically fast conformational relaxation from the minor to the dominant state in this off-pathway exchange.

A monotonous decrease of *k*_2_([L]_0_) thus can only occur for conformational selection, which has been used to identify this binding mechanism in protein systems with *k*_*e*_ *< k*_−_ ^23,24^. Conformational selection has also been identified for a system with *k*_*e*_ *> k*_−_ from an asymmetry of the two arms of the function *k*_2_ [L]_0_ left and right of the minimum ^25^. For these protein systems, only the smaller rate *k*_2_, also termed *k*_obs_, could be observed (i.e. deduced from fits of the stopped-flow relaxation curves) at the different ligand concentrations [L]_0_.

Induced fit, in contrast, is in general more difficult to identify based on stopped-flow mixing experiments, because a (near) symmetry of the function *k*_2_([L]_0_) can occur also for conformational selection. Deducing induced fit from stopped-flow experiments therefore requires a closer look at the rate constants of the induced-fit and conformational-selection models that are obtained from fitting these models to the stopped-flow data. In this model fitting, the induced-fit and conformational-selection mechanisms are simplified from the 4-state models of Fig. 2 to the 3-state models in Fig. 3 to reduce the number of fit parameters. The rates *k*_1_ and *k*_2_ are not affected by the additional “ off-pathway” conformational change in the 4-state models if the sum of the rates for this exchange is larger than *k*_1_ and *k*_2_, which is plausible for a typically fast conformational relaxation from the minor to the dominant state in this off-pathway exchange.

## Induced fit of the neurotensin receptor 1 inferred from stopped-flow mixing and saturation transfer difference NMR experiments

The neurotensin receptor 1 (NTS1) is a GPCR that is activated by the endogenous peptide neurotensin as ligand ^26^. NTS1 is primarily expressed in the central nervous system and gastrointestinal tract ^27^ and regulates neurological processes including dopamine transmission and GABAergic system modulation ^28^. For a ther-mostabilized NTS1 variant solubilized in detergent as replacement for the native membrane environment, stopped-flow mixing experiments allowed to determine the two binding relaxation rates *k*_1_ and *k*_2_ at concentrations [L]_0_ of the ligand neurotensin between 0.5 and 2.5 µM and the relaxation rate *k*_1_ at additional concentrations [L]_0_ between 3.75 and 15 µM ^1^ (see data points in Fig. 4). The protein concentration in all mixing experiments was [P]_0_ = 1 µM. The stopped-flow mixing experiments were conducted in the absence of the associated G protein, because the G protein did not induce any further changes to the conformational dynamics of the NTS1-neurotensin complex in NMR experiments ^1^. Neurotensin alone thus appears to sufficiently stabilize the active conformation of thermostabilized NTS1.

**Fig. 4.**
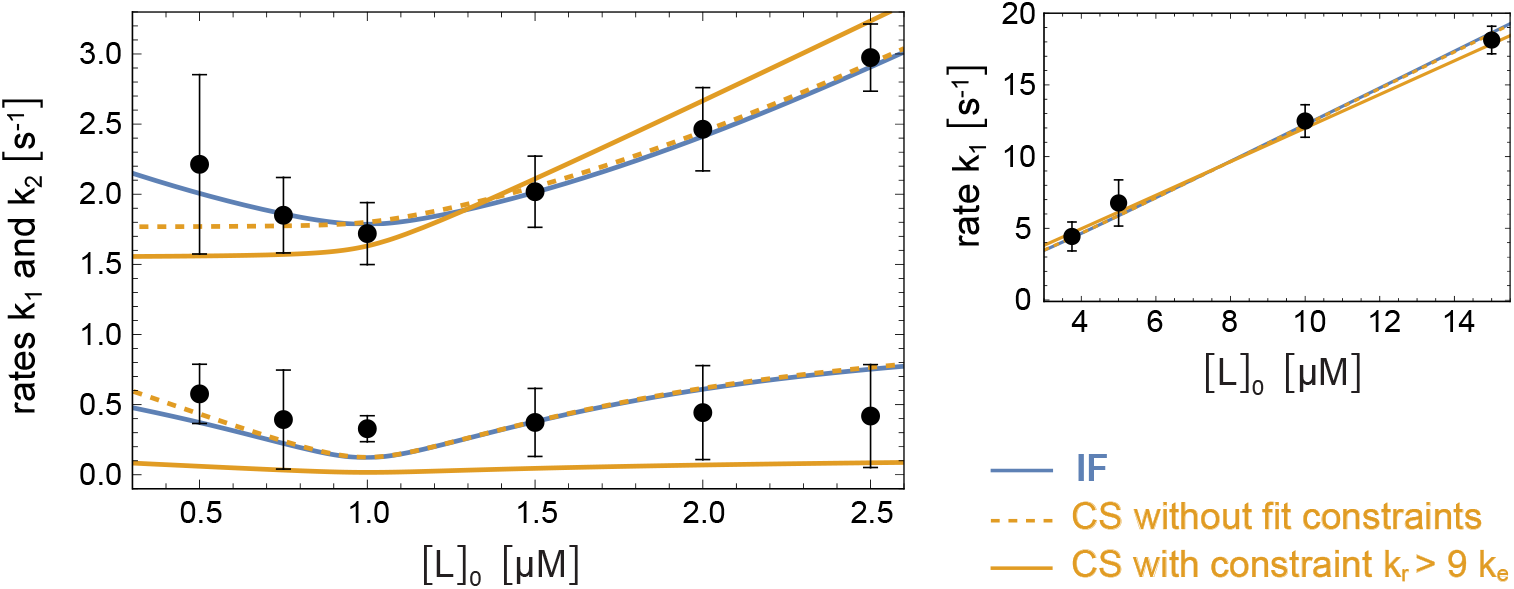
Fits of the relaxation rates *k*_1_ and *k*_2_ from stopped-flow mixing experiments of the thermostabilized receptor NTS1 and the peptide ligand neurotensin (data points) with the induced-fit (IF) and conformational-selection (CS) models (coloured lines) 1. For neurotensin concentrations [L]_0_ between 0.5 and 2.5 µM, both rates *k*_1_ and *k*_2_ could be determined from double-exponential fits of the stopped-flow relaxation curves. For neurotensin concentrations [L]_0_ between 3.75 and 15 µM, the rate *k*_1_ was determined from single-exponential fits of the initial relaxation in the stopped-flow experiments. The NTS1 concentration in all mixing experiments was [P]_0_ = 1 µM. All data points were jointly fitted with both the induced-fit and conformational-selection with the three rate constants *k*_*e*_, *k*_*r*_, and *k*_−_ as fit parameters. The fourth rate constant of the models, *k*_+_ was replaced by the experimentally measured dissociation constant *K*_*D*_ = 6 ± 2 nM. The resulting fit values are shown in Table 1. For the conformational-selection model, both an unconstrained fit and a fit with the constraints *k*_*r*_ *>* 9*k*_*e*_ was performed. In this constraint fit, the relative probability of R_2_ in the unbound state is limited to plausible values smaller than 10% (see text). The figure is adapted from Ref. 1.

To deduce the binding mechanism from the stopped-flow data, the functions *k*_1_([L]_0_) and *k*_2_([L]_0_) for the induced-fit and conformational-selection model were fitted to the data with the three rate constants *k*_*e*_, *k*_*r*_, and *k*_−_ as fit parameters (see coloured lines in in Fig. 4). The fourth rate constant of the models, *k*_+_, was replaced by experimentally measured dissociation constant *K*_*D*_ = 6 ± 2 nM of the NTS1-neurotensin complex, which depends on the rate constants of the models (see caption of Fig. 3). In the induced-fit model, *k*_*e*_ and *k*_*r*_ are the conformational transition rates between R_1_L and R_2_L, and *k*_−_ is the dissociation rate of R_1_L. In the conformational-selection model, *k*_*e*_ and *k*_*r*_ denote the transition rates between R_1_ and R_2_, and *k*_−_ is the dissociation rate of R_2_L.

**Table 1.** Values of rate constants for the induced-fit (IF) and conformational-selection (CS) model obtained from the fits in Fig. 4 in units of *s*^−1^. Number in brackets indicate 95% confidence intervals.

| fitted model | $k_e$ | $k_r$ | $k_-$ |
| --- | --- | --- | --- |
| IF | 0.015 [0, 0.04] | 1.1 [0.7, 1.6] | 0.6 [0.25, 0.95] |
| CS without constraints | 1.2 [0.7, 1.7] | 0.6 [0.2, 1.0] | 0.005 [0.004, 0.009] |
| CS with constraint $k_r > 9k_e$ | 0.16 [0, 0.8] | 1.4 [0, 0.8] | 0.001 [0, 0.004] |

Fitting the conformational-selection model without constraints on rate parameters leads to a large probability *P*(R_2_) = *k*_*e*_*/*(*k*_*e*_ + *k*_*r*_) *>* 50 % of the active conformation R_2_ in the unbound state (see Table 1), in contradiction to structural data for NTS1^29^ and the thermostabilized NTS1 variant ^30^. Fits in which *P*(R_2_) is constrained to plausible values < 10% (i.e. to rate parameters *k*_*r*_ *>* 9*k*_*e*_) poorly match the stopped-flow data (see Fig. 4). With the induced-fit model, in contrast, the stopped-flow data can be well fitted with plausible excitation and relaxation rate constants *k*_*e*_ and *k*_*r*_ for the conformational exchange between R_1_L and R_2_L, and with a dissociation rate constant *k*_−_ of the bound excited state R_1_L of about 0.6 s^−1^. Moreover, the conformational exchange rate constants *k*_*e*_ and *k*_*r*_ obtained from the stopped-flow data in the induced-fit model are in good agreement with the exchange rate constants 0.08 *s*^−1^ and 1.23 *s*^−1^ measured in saturation transfer difference (STD) NMR experiments of NTS1 bound to a fluorinated neurotensin variant. This agreement of conformational exchange rates in the bound state obtained from distinct experiments is a rather strong indication of induced fit as the binding mechanism.

## Induced fit of the β_2_-andrenergic receptor inferred from a decrease of ligand association rates after stabilization of the active receptor conformation

The β_2_-andrenergic receptor (β_2_AR) is a prototypic GPCR that recognizes epinephrine (adrenaline) as ligand and mediates a variety of physiological responses including smooth muscle relaxation and bronchodilation ^31^. To investigate the interplay between ligand binding on the extracellular side and G-protein binding on the intracellular side of β_2_AR, Devree et al. ^2^ monitored the association kinetics of ligands to β_2_AR-G protein complexes. In these complexes, the G protein is either bound to the nucleotide GDP, or nucleotide-free.

For the nucleotide-free G protein in complex with β_2_AR, Devree et al. ^2^ observed significantly reduced ligand association rates compared to the GDP-bound G protein. A significant reduction of ligand association rates was also observed for β_2_AR bound to the nanobody Nb80, which stabilizes the active β_2_AR conformation ^32^. From the similar effect of Nb80 and nucleotide-free G protein on ligand association rates, Devree et al. ^2^ concluded that they both stabilize the active β_2_AR conformation, and that ligand binding is impaired in this conformation.

In the kinetic mechanism for β_2_AR activation suggested by Devree et al. ^2^ (see Fig. 5), the nucleotide-free G protein is associated to the active β_2_AR conformation, while the GDP-bound G protein is associated to the inactive β_2_AR conformation, which is the ligand-binding-competent conformation in this mechanism. The significantly reduced ligand association rates of β_2_AR in complex with the nucleotide-free G protein can be understood from the stabilization of the active conformation in this complex. This kinetic mechanism is the induced-fit mechanism of Fig. 2 with R_1_ corresponding to the inactive β_2_AR conformation in complex with the GDP-bound G protein, and R_2_ corresponding to the active β_2_AR conformation in complex with the nucleotide-free G protein.

**Fig. 5.**
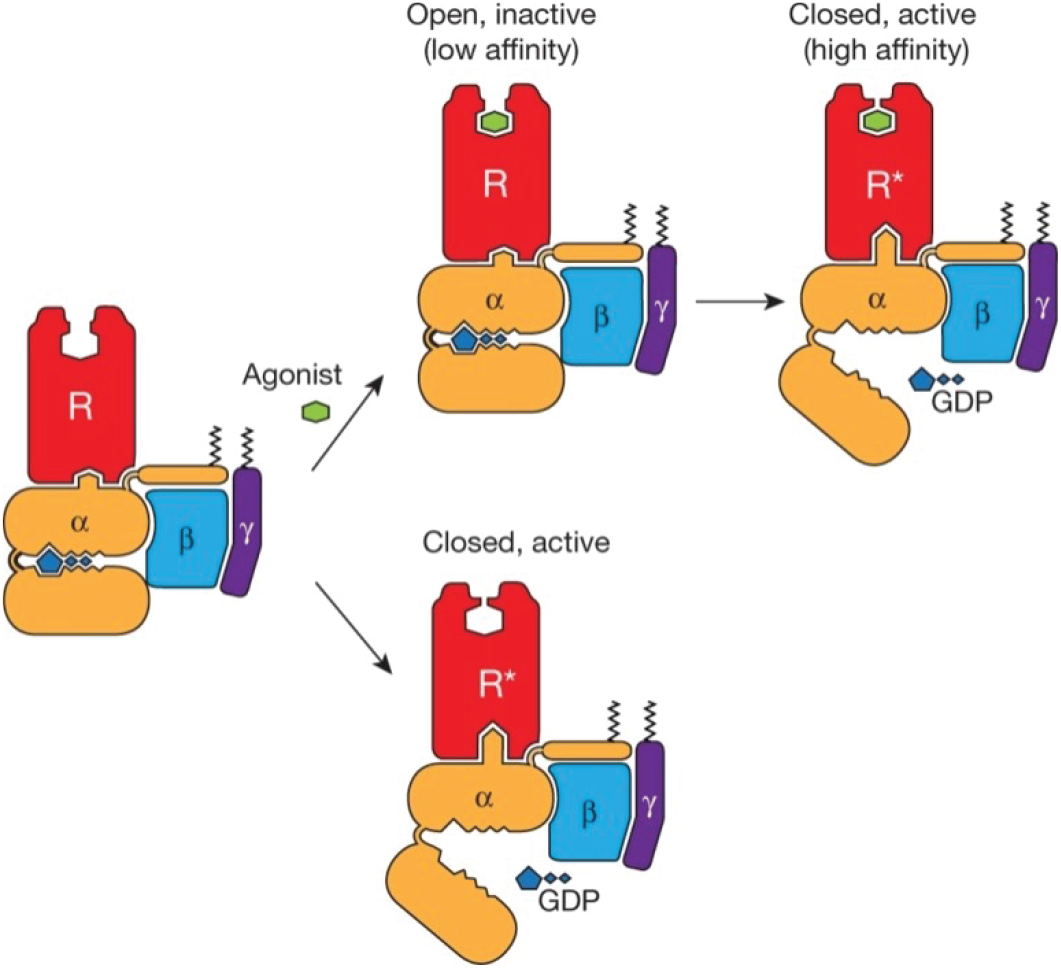
Kinetic mechanism for β_2_AR activation 2. The G protein is represented as heterotrimer with subunits α, β, and γ. The G protein with GDP-bound α-subunit is associated with the inactive, open β_2_AR, and the nucleotide-free G protein is associated with the active, closed β_2_AR conformation. Binding of ligand agonists only occurs in the inactive β_2_AR conformation in this mechanism. The figure is reproduced with permission from Ref. 2.

As structural explanation for impaired ligand binding in the active β_2_AR conformation, Devree et al. ^2^ point out a lid-like structure formed by two aromatic residues that closes over the ligandbinding site in this conformation. Mutating one of these bulky aromatic residues to the small residue alanine diminishes the effect of the nanobody Nb80 on ligand association rates, which supports the lidding effect of these residues ^2^.

## Induced fit as likely allosteric mechanism of small-molecule-activated and peptide-activated GPCRs

Constriction and closing of the ligand-binding site in the active conformation is rather common for small-molecule-activated and peptide-activated GPCRs. Similar to β_2_AR, the active structure of the muscarinic acetylcholine receptor exhibits a lid-like structure over the ligand binding site ^2,33^. For this receptor as well as the µ-opioid receptor, Devree et al. ^2^ reported a decrease of ligand association rates after stabilization of the active conformation akin to β_2_AR, which indicates that the induced-fit allosteric mechanism of Fig. 5 is shared by the receptors. Also for the P2Y12 receptor, a lid-like structure over the ligand binding site has been observed for a complex with a close analogue of the endogenous ligand ADP ^34^.

Compared to small-molecule-activated GPCRs, the binding sites of peptide-activated GPCRs tend to be larger and more open to accommodate the peptide ligands. However, structural data indicate a constriction of the binding site in the active conformation leading to tight interactions between receptor and peptide ^29,35–38^. This binding-site constriction provides a structural explanation for the induced-fit binding of NTS1 to neurotensin ^1^, and lends plausibility to induced fit as general allosteric mechanism of peptide-activated GPCRs.

## Acknowledgements

T.R.W. thanks the Max Planck Society for funding.

